# CaptureCompendium: a comprehensive toolkit for 3C analysis

**DOI:** 10.1101/2020.02.17.952572

**Authors:** Jelena M. Telenius, Damien J. Downes, Martin Sergeant, A. Marieke Oudelaar, Simon McGowan, Jon Kerry, Lars L.P. Hanssen, Ron Schwessinger, Chris Q. Eijsbouts, James O.J. Davies, Stephen Taylor, Jim R. Hughes

## Abstract

DNA folding within nuclei is a highly ordered process, with implications for gene regulation and development. An array of chromosome conformation capture (3C) methods have been developed to investigate how DNA is packaged within nuclei and to interrogate specific interactions. While these methods use different approaches to examine target loci (many-versus-all) or the entire genome (all-versus-all), they all rely on the core principle of endonuclease digestion and proximity-based ligation to re-arrange genomic order to reflect the three-dimensional nuclear conformation. This sequence reorganization creates novel chimeric DNA fragments which require specialist bioinformatic tools to analyze and visualize. Despite this need for specialist bioinformatic skills, the core biological importance of genome folding has seen widespread methodological uptake. To service the needs of experimentalists using the many-versus-all Capture-C family of methods we have developed CaptureCompendium; a toolkit of software to simplify the design, analysis and presentation of 3C experiments.

## Introduction

The arrangement of the genome into the nucleus is highly regulated to facilitate correct interactions between promoters and their requisite enhancers. Chromosome conformation capture (3C) has emerged as a highly malleable method for studying both local and global trends in genome folding^1^. All 3C methods rely on the underlying principle that following DNA digestion, generally with a restriction endonuclease (RE), fragment ends in proximity will ligate together at high frequency. This digestion and proximity driven ligation functions to re-organize the linear genome sequence to reflect its 3D structure^2^. This re-organized sequence is then examined primarily by targeted high-resolution many-versus-all approaches, e.g. NG Capture-C^3^, 4C-seq^4^, UMI-4C^5^, i4C-seq^6^, and Tri-C ^7^; or lower resolution approaches across the genome, e.g. HiC^8^ and Capture-HiC^9^.

The importance of genome folding to transcriptional regulation has brought about the application of 3C methods to investigate a range of questions, including basic biological problems and the identification of disease relevant gene-to-polymorphism interactions^10,11^. Owing to the enhanced signal provided by targeted sequencing, oligonucleotide capture methods, based on Capture-C^3,7,9,12–14^, are emerging as a preferred method to look at loci of interest. A standard Capture-C experiment consists of three main bioinformatic components: target selection and oligonucleotide probe design; sequence mapping with read classification; and statistical comparison with data presentation. For each of these processes an array of software tools is available, including HiCapTools^15^, GOPHER^16^, Capsequm^12^, capC-MAP^17^, FourCSeq^18^, peakC^19^, CHiCAGO^20^, peaky^21^, ChiCMaxima^22^, HiC-Pro^23^, and 4See^24^; yet, to date no comprehensive pipeline for the complete design, mapping and interpretation of 3C experiments has been made available. This lack of an easily implemented, automated workflow for Capture-C based experiments forms a barrier to the general use of such methods, particularly in non-specialist groups. To address this need we have developed the CaptureCompendium toolkit (Fig. 1) which combines oligonucleotide design (Capsequm2), sequence mapping and data extraction (CCseqBasic), and statistical analyses with data presentation and distribution (CaptureCompare, Tri-C, CaptureSee). This workflow streamlines the bioinformatic process required to generate high quality 3C data using Capture-C experiments.

**Figure 1.**
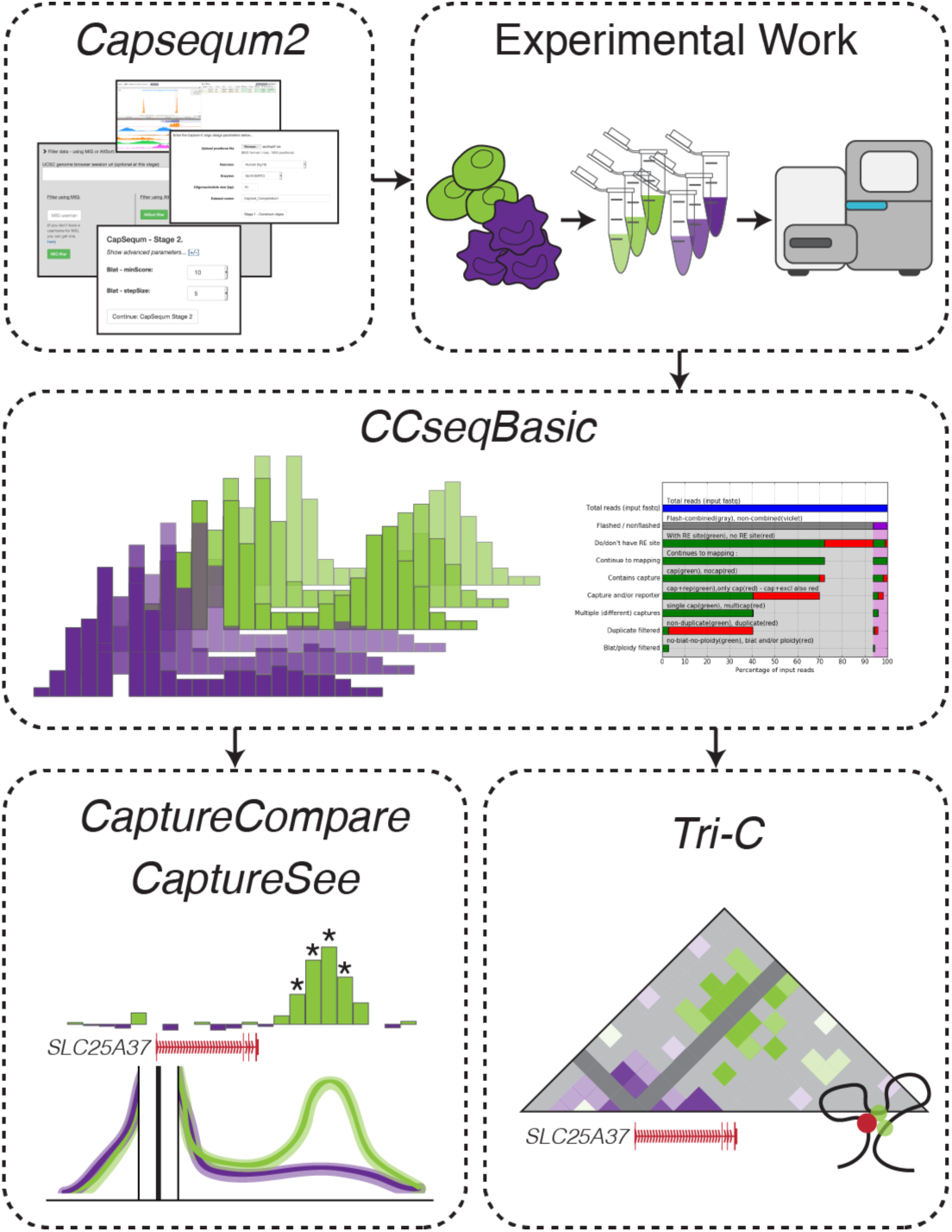
Overview of CaptureCompendium. Biotinylated oligonucleotides for 3C experiments can be designed using Capsequm2. Following the experimental work, individual samples are analysed and QC is performed in CCseqBasic. Whole experiments, with replicates and comparators are analysed in CaptureCompare and visualized in CaptureSee. Multi-interaction hubs in Tri-C can also be detected using CCseqBasic.

## RESULTS

### Capsequm2: A user-friendly interface for probe design

Targeted enrichment with biotinylated DNA oligonucleotides forms the basic tenet of Capture-C methods^3^. Probe design is essential to a successful experiment; probes must be situated near to a RE cut site, be proximal to the object of interest, and not be located on a highly repetitive element. These parameters allow for enrichment of informative on-target chimeric sequencing fragments, while simultaneously avoiding off-target sequencing. Currently, we know of three tools are available for probe design: HiCapTools - a command line-based method^15^, GOPHER – an installable JAVA package^16^, and Capsequm – an online interface^12^. As a webtool, Capsequm provides the most user-friendly option for initial probe design but previously required integration with MIG^25^, a registration-based service, for probe filtering.

We redeveloped Capsequm (http://apps.molbiol.ox.ac.uk/CaptureC/cgi-bin/CapSequm.cgi) with a web-based filtering portal (AltSort) to generate Capsequm2 (Fig. 2). Capsequm2 allows researchers to select from a range of 11 genomes covering seven species and the five commonly used RE, as well as allowing a choice in probe length for design to targets of interest. Following fragment identification and BLAT processing, probes can be filtered using AltSort. In AltSort probes filtering are automatically categorised as good or bad using recommended settings for duplication number, BLAT density score (measuring likely off-target capture) (Kent 2002), G/C content, CpG content and, repeat content (Table 1). Probes can be visually inspected with integrated UCSC browser tracks, and the default parameters adjusted for all or specific regions of interest. Upon selection of desired probes, users can then download the oligonucleotide sequences, ready for ordering. This download also provides correctly formatted viewpoint input files required for subsequent analysis with the downstream tools CCseqBasic and CaptureCompare.

**Table 1.** Default AltSort parameters for probe design.

| Filter | “Good” | “Bad” |
| --- | --- | --- |
| Duplicates | $\leq 2^*$ | $\geq 3$ |
| Density | $\leq 40$ | $> 40$ |
| % G/C content | $\leq 60$ | $\geq 60$ |
| Repeat | False | True |

\*Often interactions at duplicated genes, e.g. *Hba-1*, *Hba-2*, can still be understood.

**Figure 2.**
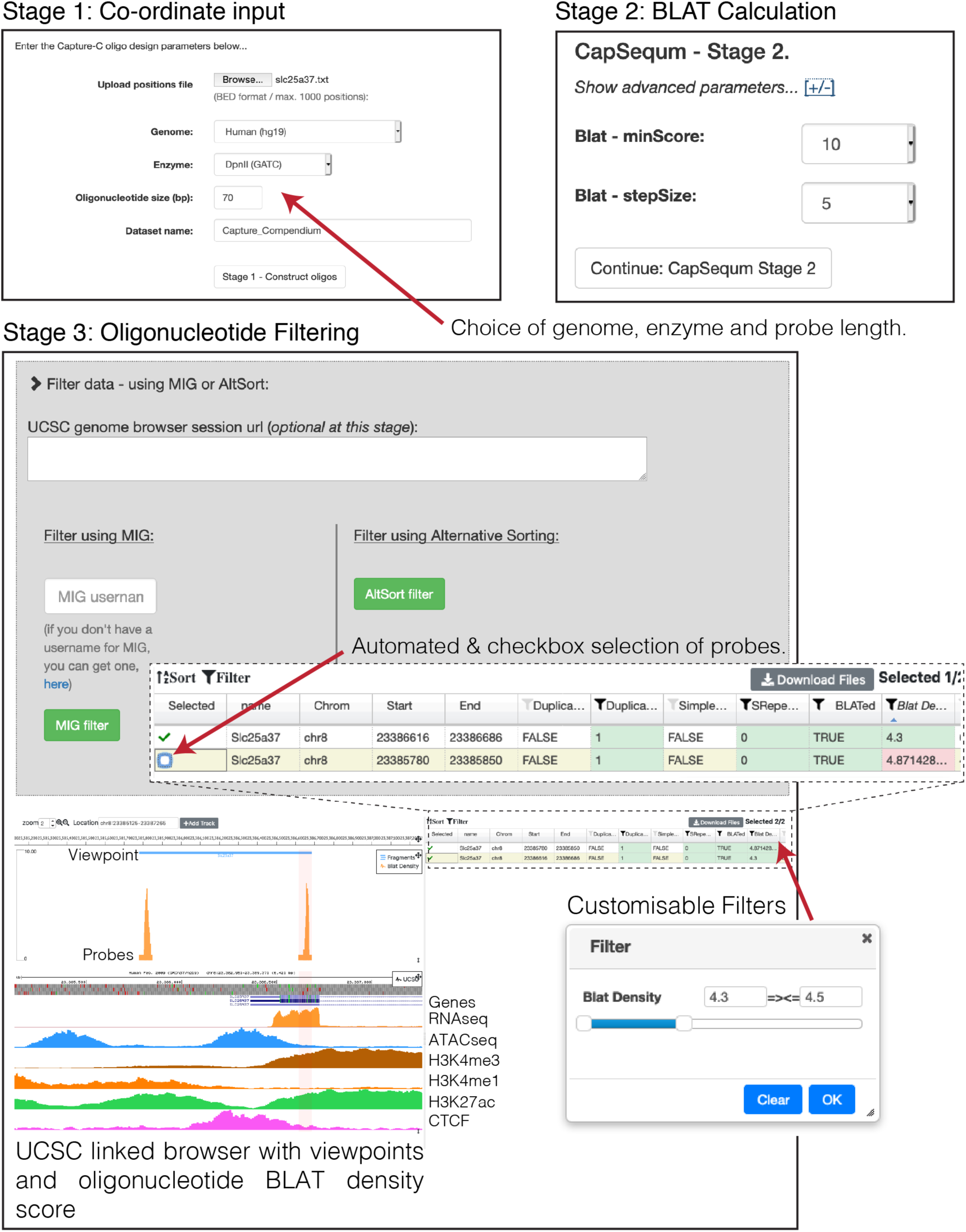
CapSequm2 provides a user-friendly web interface for oligonucleotide design. The online tool allows design to desired capture locations with selection of genome from Human (hg18/19/38), Mouse (mm9, mm10), Chimpanzee (panTro3), Zebrafish (danRer7, danRer10), *Drosophila melanogaster* (dm3), Chicken (galGal4, galGal5), & Sickleback (BROADS1), of enzyme (*Dpn*II, *Nla*III, *Bam*HI, *Bgl*II, *Hind*III), and of oligonucleotide size. Designed oligonucleotides are then quality filtered using MIG or AltSort which allows integration with chromatin data. AltSort automatically selects oligonucleotides on recommended BLAT density, duplication, and repeat values which can be manually adjusted.

### CCseqBasic: Data extraction and quality control

Analysis of sequencing data with the CCseqBasic command line tool (Fig. 3a) is launched using the probe design parameter file provided by AltSort. Enriched chimeric fragments from the Capture-C family of methods, NG Capture-C, LI-Capture-C, Tag Capture-C, Tri-C, are generally sequenced using 150-bp paired-end reads^3,7,13,26^; this gives 300-bp of total sequence, therefore the majority of read-pairs overlap and the chimeric fragment can be reconstructed through flashing^27^. RE sites in chimeric fragments generated from 3C digestion and ligation are then identified in both flashed and non-flashed reads and digested-*in silico*. These digested reads are then mapped and categorised as capture, occurring within a targeted RE fragment, or reporter, occurring in any other fragment. Because Capture-C methods use sonication or tagmentation to generate random ends for reads, they can then be PCR duplicate filtered. All of these steps are carried out automatically by CCseqBasic with minimal user input. The main output of CCseqBasic is a convenient UCSC-loadable data hub which contains reporter tracks for each viewpoint (Fig. 4), complete with sample quality control metrics for 3C library quality and enrichment efficiency (Fig. 3b), as well as sequence quality control in the form of fastQC reports. Additionally, all generated data is stored in an easy-to-follow hierarchical folder structure allowing re-analysis, or immediate further processing with CaptureCompare or Tri-C.

**Figure 3.**
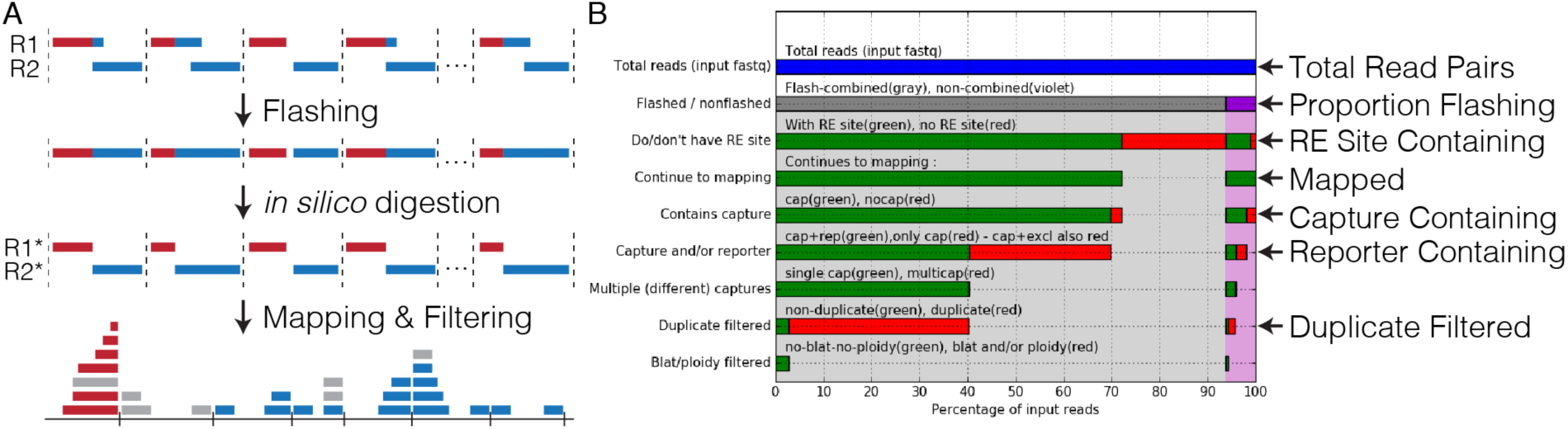
CCseqBasic extracts informative data from 3C sequence reads. **(a)** 3C libraries are analysed by reconstructing original fragments by flashing. Subsequent *in silico* digestion of reads resolves chimeric fragments, and allows mapping with bowtie. Mapped reads are identified as “Capture” (red): coming from a targeted viewpoint, or “Reporter” (blue): coming from an interacting fragment. Capture-reporter read pairs are then filtered for PCR duplicates and off target capture sites (grey). **(b)** 3C library and capture quality metrics are reportable as fractions of total reads to allow easy tracking of experimental success, or where necessary, facilitate troubleshooting.

**Figure 4.**
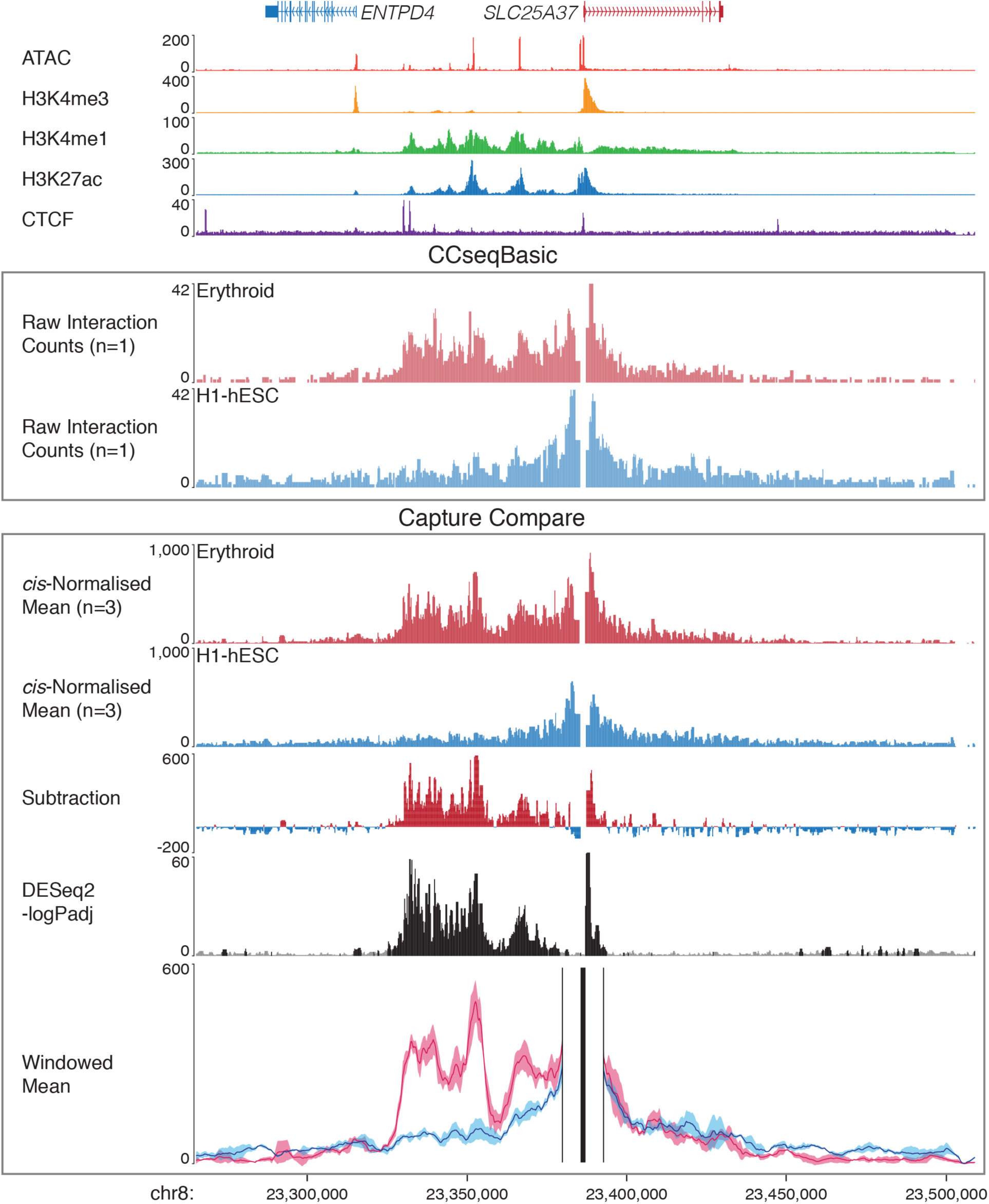
CaptureCompare allows statistical analysis of experiments. The outputs of CCseqBasic and CaptureComparecan be directly loaded into UCSC for interpretation with chromatin epigenetic data. Example ouput is a Capture-C experiment from the viewpoint of the human mitoferrin gene *SLC25A37*. Output tracks include raw reporter counts, *cis*-normalised reporter counts, subtraction between comparison samples, and -logPadj values from DESeq2 on a per fragment basis. A pdf of windowed mean interactions with standard deviations is also provided. For DESeq2, significantly different fragments have been coloured black (FDR q<0.05).

### Tri-C: Investigation of 3-way interaction hubs

Most 3C methods interrogate single, pairwise interactions to generate a population view of interaction frequencies. Recently, interest has been given to the formation of multi-way interaction hubs^28–30^. By enriching for chimeric fragments containing multiple ligation junctions, Tri-C allows investigation of these multi-way interactions from single-alleles^7^. Tri-C interaction matrices are generated using an optional flag, which can be implemented during either the initial run or re-analysis with CCseqBasic. When using Tri-C analysis, users can specify the genomic regions to analyse, sonication fragment size, and bin size to match their specific experiment. The output of this analysis is a pdf of an interaction matrix for the region surrounding the viewpoint of interest (Fig 5.).

**Figure 5.**
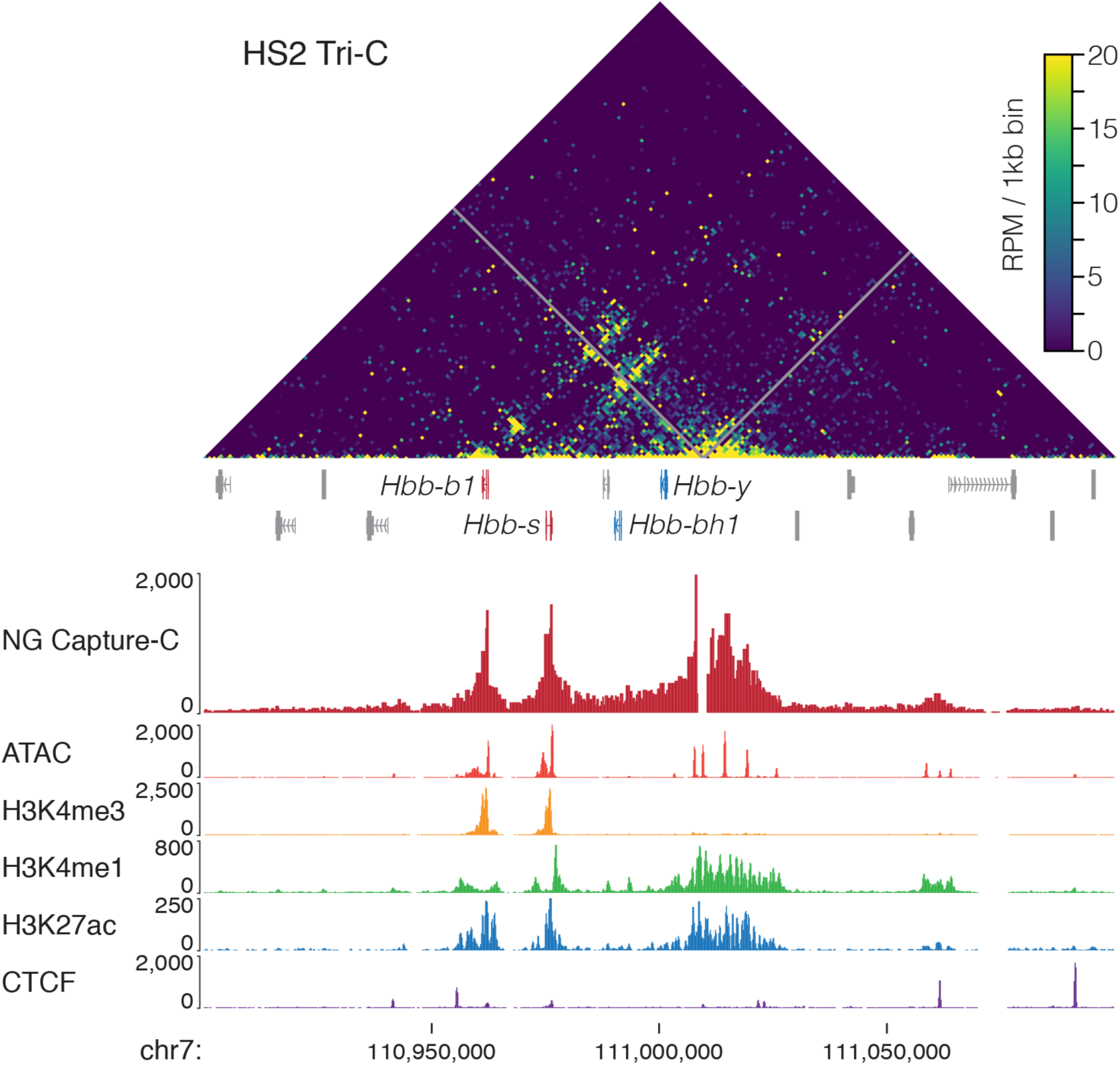
Detection of multi-way interaction hubs with Tri-C analysis. The output interaction map from Tri-C analysis of the viewpoint from the HS2 element of the mouse beta-globin Locus Control Region (LCR) shows concurrent interaction between HS2, the active beta-globin genes, and other elements within the LCR.

### CaptureCompare: Statistical analysis and sample comparison

A major strength of Capture-C methods is their ability to confidently distinguish PCR duplicates, while simultaneously generating data from multiple replicates and cell types by multiplexing samples^3^. This allows experiments to be designed with advanced statistical analysis in mind, such as the comparison between cell types, differentiation states, genetic models, drug treatment or any other biological question of interest. As with analysis by CCseqBasic, CaptureCompare is implemented using a command line interface and AltSort-generated parameter file. CaptureCompare extracts the raw interaction count data for each viewpoint from replicates of multiple conditions and generates a UCSC loadable hub containing normalised mean interactions, standard deviation, and difference in interaction level for different conditions on a per RE fragment basis (Fig. 4). Additionally, CaptureCompare generates and implements a modifiable R script for DESeq2 analysis^31^, which compares RE fragments to determine if observed differences between conditions reach significance. CaptureCompare also automates the preparation of the necessary files for more complex interaction frequency analysis via the Bayesian modelling tool peaky^21^. Finally, to facilitate presentation, a pdf is generated for each viewpoint with publication-style windowed interaction profiles (Fig. 4).

### CaptureSee: A simple interface for navigating and sharing experiments

On-going methodological improvements have increased the ability of Capture-C methods to target thousands rather than tens to hundreds of loci. Exploration of these large datasets is difficult when using the UCSC browser. To facilitate easy interaction with 3C data, and provide a platform for sharing results, we developed the web-based platform CaptureSee. CaptureSee takes the output hub from CaptureCompare to generate a browsable repository of viewpoints in both per RE fragment and windowed formats (Fig 6.). Within CaptureSee users can quickly jump between capture viewpoints in the genome, to see both the total number and percent of *cis* reporter counts for each viewpoint on a per sample basis, and filter DESeq2 results to find significantly interacting RE fragments.

**Figure 6.**
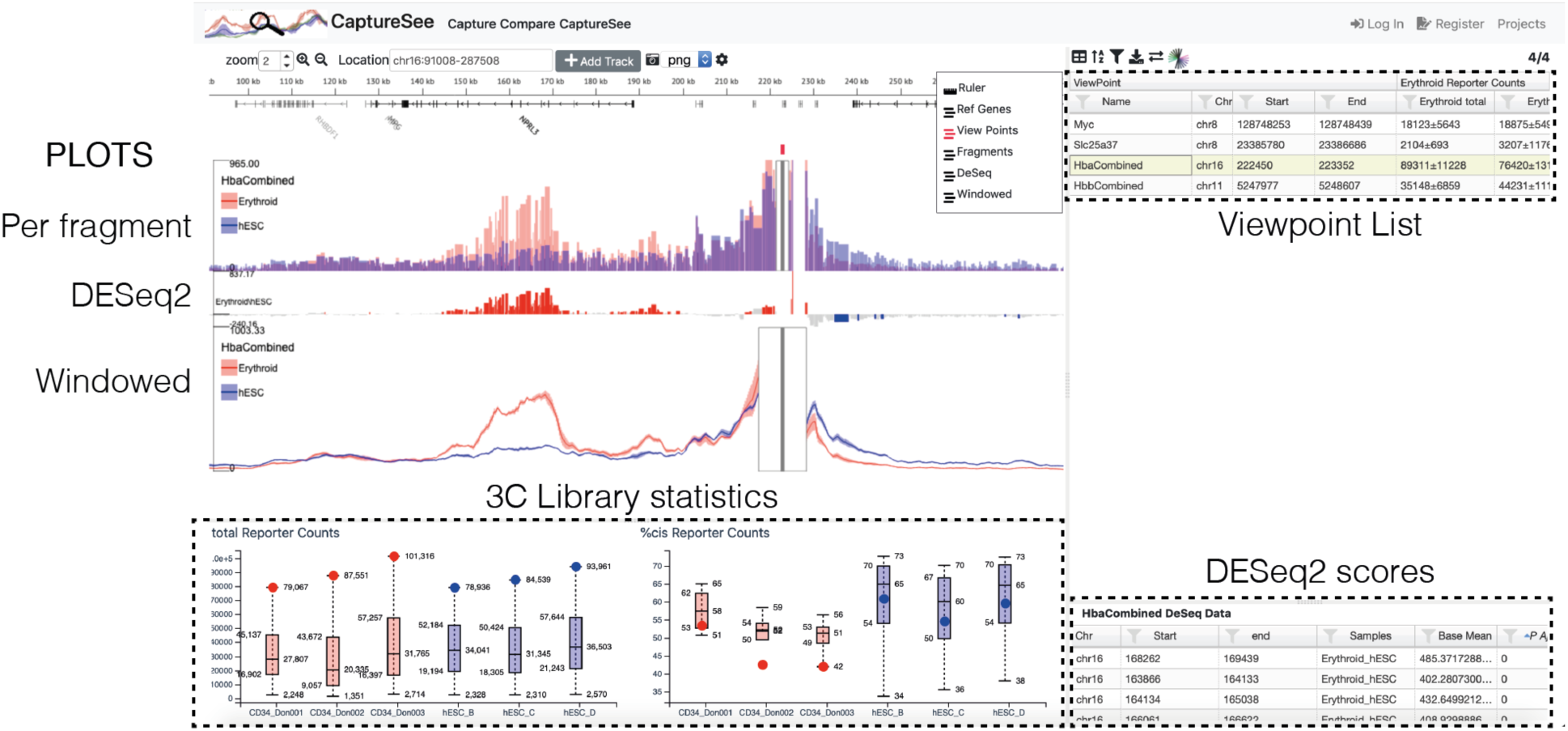
Data exploration and sharing with CaptureSee. Data is loaded into CaptureSee using the CaptureCompare generated public hub. Within CaptureSee users can quickly move between viewpoints; with both per fragment and widowed views. In addition to 3C profiles, reporter counts and percent *cis* interactions are provided for each sample 3C library. DESeq2 logPadj scores are provided on a per fragment basis and can be filtered for level of significance.

## DISCUSSION

The importance of genome folding for gene regulation has come to the fore in recent years. An extensive array of techniques in fixed and native cells has been developed to examine this folding based on the premise of proximity ligation. Despite being based on a simple premise, 3C sequencing generates a complex bioinformatic problem of mapping and interpreting chimeric DNA fragments. To meet this need, an equally wide range of tools have been developed to either design experiments^12,15,16^, map sequence reads^3,17^, or interpret interaction profiles^18–22^. While some of these tools have been developed with a non-bioinformatically trained user in mind, many are complex to implement. The absence of a coherent analysis workflow necessitates the definition and testing of appropriate solutions and tools for each step of design, analysis, interpretation and visualisation which must then be assembled into a functioning bioinformatics pipeline, before such experiments can be productively undertaken. This has slowed the general uptake into other fields, such as genetics, clinical studies and developmental biology, though Capture-C methods have proven utility in these areas^10,32–41^.

We have generated a systematic, semi-automated workflow for the design and analysis of Capture-C based 3C experiments. This pipeline is constructed with the end-user in mind, including easy to navigate web tools where computationally possible. For the command line tools the default settings will suffice for general use, and user input is only required at three stages. More advanced users will find their needs met, with options and flags available at all stages to customise analysis – such as the choice in mapping algorithm or using SNPs to produce allele-specific profiles^3^. Furthermore, the availability of well-documented open source software in commonly used programming languages (Unix, perl, R) means further modifications can be easily incorporated. Therefore, CaptureCompendium represents a significant advance for the implementation of capture-based 3C methodologies that will enable widespread use of these powerful techniques to explore genomes structure.

## METHODS

Capture-C data presented in this work are available on GEO (GSE129378) and can be visualised at (http://capturesee.molbiol.ox.ac.uk/projects/capture_compare/1387). The Tri-C interaction profile was generated from published data^7^ (GSE107755). CaptureCompendium with links to software, installation guide, usage manual and tutorial vignettes is available online (http://userweb.molbiol.ox.ac.uk/public/telenius/CaptureCompendium/). Capsequm2, AltSort and CaptureSee are available as web tools with full descriptions on the websites (http://apps.molbiol.ox.ac.uk/CaptureC/cgi-bin/CapSequm.cgi and https://capturesee.molbiol.ox.ac.uk/). To facilitate saving and sharing data, CaptureSee requires users to register for a free account. CCseqBasic is modified from previous published versions^3^, and now contains Tri-C. CCseqBasic is available on GitHub (https://github.com/Hughes-Genome-Group/CCseqBasicS) and implements a suite of tools to analyse data. Briefly adaptor sequences a trimmed with TrimGalore (https://www.bioinformatics.babraham.ac.uk/projects/trim_galore/), reads are flashed using FLASH^27^, then *in silico* digested, and mapped with either bowtie or bowtie2^42,43^. Read quality is measured at multiple stages of processing using FastQC (http://www.bioinformatics.babraham.ac.uk/projects/fastqc/). Reads are classified as capture or reporter depending on mapping location and PCR duplicates are filtered based on duplication of random end pairs generated by sonication. Skew arising from co-capture from oligonucleotides at two independent viewpoints is excluded by masking all viewpoints included within a Capture-C experiment. CaptureCompare is available on github (https://github.com/Hughes-Genome-Group/CaptureCompare), and uses custom perl and R scripts to analyse each viewpoint independently by normalising reads per 100,000 *cis* interactions, combining replicates using BEDtools^44^, generating mean and subtraction data and inputting read counts into DESeq2^31^ treating each restriction fragment as an individual object. Bedgraphs are converted to bigwigs with deepTools2^45^. Windowed plots are generated with adjustable bin and window sizes using custom R scripts.

## Acknowledgements

This work was carried out as part of the WIGWAM Consortium (Wellcome Investigation of Genome Wide Association Mechanisms) funded by a Wellcome Trust Strategic Award (106130/Z/14/Z) and Medical Research Council (MRC) Core Funding (MC_UU_12009). We wish to thank all the testers and users of the tools, particularly Duantida Songdej and Nigel Roberts, for dedicated and detailed error reporting and alpha testing, and Jason Torres and Gabriele Mingiolini for portability testing. We acknowledge the CCB computational cluster and the AVI research group for providing the computational resources and system administration for the software development and production runs. Wellcome Trust Doctoral Programmes supported R.S. (203728/Z/16/Z), C.Q.E. (203141/Z/16/Z), A.M.O. (105281/Z/14/Z) and L.L.P.H (099684/Z/12/Z). A.M.O was also supported by the Stevenson Junior Research Fellowship (University College, Oxford). J.O.J.D. is funded by an MRC Clinician Scientist Award (MR/R008108) and received Wellcome Trust Support (098931/Z/12/Z).

